# De novo profile generation based on sequence context specificity with the long short-term memory network

**DOI:** 10.1101/240515

**Authors:** Kazunori D Yamada, Kengo Kinoshita

## Abstract

Long short-term memory (LSTM) is one of the most attractive deep learning methods to learn time series or contexts of input data. Increasing studies, including biological sequence analyses in bioinformatics, utilize this architecture. Amino acid sequence profiles are widely used for bioinformatics studies, such as sequence similarity searches, multiple alignments, and evolutionary analyses. Currently, many biological sequences are becoming available, and the rapidly increasing amount of sequence data emphasizes the importance of scalable generators of amino acid sequence profiles. We employed the LSTM network and developed a novel profile generator to construct profiles without any assumptions, except for input sequence context. Our method could generate better profiles than existing de novo profile generators, including CSBuild and RPS-BLAST, on the basis of profile-sequence similarity search performance with linear calculation costs against input sequence size. In addition, we analyzed the effects of the memory power of LSTM and found that LSTM had high potential power to detect long-range interactions between amino acids, as in the case of beta-strand formation, which has been a difficult problem in protein bioinformatics using sequence information. We demonstrated the importance of sequence context and the feasibility of LSTM on biological sequence analyses. Our results demonstrated the effectiveness of memories in LSTM and showed that our de novo profile generator, SPBuild, achieved higher performance than that of existing methods for profile prediction of beta-strands, where long-range interactions of amino acids are important and are known to be difficult for the existing window-based prediction methods. Our findings will be useful for the development of other prediction methods related to biological sequences by machine learning methods.

## INTRODUCTION

Amino acid sequence profiles or position-specific scoring matrices (PSSMs) are matrices in which each row contains evolutionary information regarding each site of a sequence. PSSMs have been widely used for bioinformatics studies, including sequence similarity searches, multiple sequence alignments, and evolutionary analyses. In addition, modern sequence-based prediction methods of protein properties by machine learning algorithms often use PSSMs derived from input sequences as input vectors of the prediction. A PSSM is typically constructed from multiple sequence alignment obtained by a similarity search of a query sequence against a huge sequence database such as nr or UniProt [1]., and subsequently, the PSSM is refined by iterative database searches. The iteration is a type of machine learning process that improves the quality of profiles gradually. In recent years, HHBlits has been considered the most successful profile generation method [2]. HHBlits generates profiles by iterative searches of huge sequence databases, as in the case of PSI-BLAST [3]; however, HHBlits uses the hidden Markov model (HMM) profile, whereas PSI-BLAST adopts PSSM. To the best of our knowledge, these methods can produce good profiles on the basis of the performance of similarity searches, but they require an iterative search of a query sequence; therefore, the profile construction time depends on the size of the database. The recent increase in available biological sequences has made it more difficult to construct profiles.

In this context, de novo profile generators such as CSBuild [4, 5] and RPS-BLAST (DELTA-BLAST) [6] have been developed to reduce the cost of profile generation, although RPS-BLAST is not exactly a de novo profile generator because it explicitly uses an external profile database. CSBuild internally possesses a 13-mer amino acid profile library, which is a set of sequence profiles obtained by iterative searches of divergent 13-mer sequences. CSBuild searches short profiles against the short profile library for every part of a sequence and subsequently constructs a final profile for the sequence by merging the short profiles. CSBuild can reduce the profile construction time using precalculated short profiles; however, there is no theoretical evidence demonstrating that a PSSM can be constructed by integrating patchworks at the short (13-mer) sequence window. In other words, the previous study assumed that the protein sequences had a short context-specific tendency for the residues. This is also the case with RPS-BLAST, in which a batch of profiles obtained by searches of a query sequence against a precalculated profile library is assembled to construct a final profile.

Recently, neural networks have attracted increasing attention from various research areas, including bioinformatics. Neural networks are computing systems that mimic biological nervous systems of animal brains. Theoretically, if a proper activation function is set to each unit in the middle layer(s) of a network, it can approximate any function [7]. In recent years, neural networks have been vigorously applied to bioinformatics studies. In particular, deep learning algorithms are typically applied to neural networks. For example, several studies have applied deep learning algorithms to predict proteinprotein interactions [8, 9], protein structures [10, 11], residue contact maps [12], and backbone angles and solvent accessibilities [13]. The successes of deep learning algorithms have been realized by complex factors, such as recent increases in available data, improvements in the performance of semiconductors, development of optimal activation functions [14], and optimization of gradient descent methods [15]. These various factors have enabled calculations that were thought to be infeasible, and modern deep learning algorithms now not only stack the layers of multilayer perceptrons but also generate various types of inference methods, including stacked autoencoders, recurrent neural networks (RNNs), and convolutional neural networks [14].

The RNN is one of the most promising deep learning methods. More specifically, long short-term memory (LSTM) [16], an RNN, can be a judicious method for learning the time series or context of input vectors. Namely, with LSTM, it may be possible to learn an amino acid sequence context to predict the internal properties of amino acid sequences. The memory of LSTM is experimentally confirmed to be able to continue for more than 1,000 time steps, although theoretically, it can continue forever [16]. This memory power may be sufficient to learn features from protein sequences, for which lengths are generally less than 500 amino acids. In addition, compared with window-based prediction methods, we do not need to assume that some protein internal properties, such as secondary structure, steric structure, or evolutionary information, are formed in some lengths of amino acid sequences, as in the case of CSBuild, which assumes 13-mers. LSTM can even learn such optimal lengths of context automatically throughout learning. This characteristic of LSTM is thought to be more suitable for protein internal property predictions. Indeed, several machine learningbased prediction methods utilizing the LSTM network for protein property prediction have been successful applied [13, 17, 18].

In this study, we attempted to develop a de novo profile generator that mimicked the ability of the existing highest performance profile generation method, HHBlits, using an LSTM network, expecting our generator to be able to include the ability to input whole protein sequences. In addition, we analyzed the importance of sequence context in the prediction and performance of LSTM to solve specific biological problems through our computational experiments.

## METHODS

### Learning dataset

We conducted iterative searches using HHBlits version 2.0.15 with the default iteration library provided by the HHBlits developer and generated profiles of the sequences in Pfam [19], where the sequences were clustered by kClust version 1.0 [20] and the maximum percent identity for all pairs of sequences was less than 40% (Pfam40). Because we used the SCOP20 test dataset as a benchmark dataset for the performance of profile generators (see below), we excluded highly similar sequences with any sequences in the SCOP20 test dataset from the Pfam40 dataset using gapped BLAST (blastpgp) searches prior to the iterative search, where we considered retrieved sequences with an e-value of less than 10^−10^ as the highly similar sequences. The number of HHBlits iterations was set to three. Although HHBlits produces HMM profiles, we converted these profiles to PSSMs by extracting amino acid emission frequencies of match states. Finally, we set the generated profiles as target vectors and its corresponding sequences as input vectors in learning steps. Namely, in our learning scheme, each instance included an *N* dimension vector (sequence) as an input vector and a 20 × *N* dimension vector (profile) as a target vector, where *N* represents sequence length and 20 is the number of types of amino acid residues.

### Learning network

We designed a network with an LSTM layer, as shown in Figure 1a. In the learning steps, each amino residue in the input sequence was converted to a 400-dimension float vector by a word embedding method [21]. After the word embedding process, the input vectors were processed by an LSTM layer followed by a fully connected layer. The output of the network was set to a solution of the softmax function of the immediately anterior layer. We set the unit size of each gate of the LSTM unit to 3,200. As a cost function, we used the root mean square error between an output of the network and a target vector. As an optimizer of the gradient descent method, Adam was used [15]. As an LSTM unit, we utilized an extended LSTM with a forget gate [22], as shown in Figure 1b. In Figure 1b, the top, middle, and bottom sigmoid gates represented the input, forget, and output gates, respectively. For regularization, we used a dropout method against weights between an input layer and an LSTM layer with a drop ratio of 0.5. We observed learning and validation curves to avoid overfitting and stopped learning steps at 5,000 epochs. Because we could not deploy whole sequence data into the memory space in our computational environment, we randomly selected 40,000 sequences (about 1/40th of all sequences) and learned them as a one epoch. Therefore, an epoch in this study was about 40 times the typical epoch.

**Figure 1.**
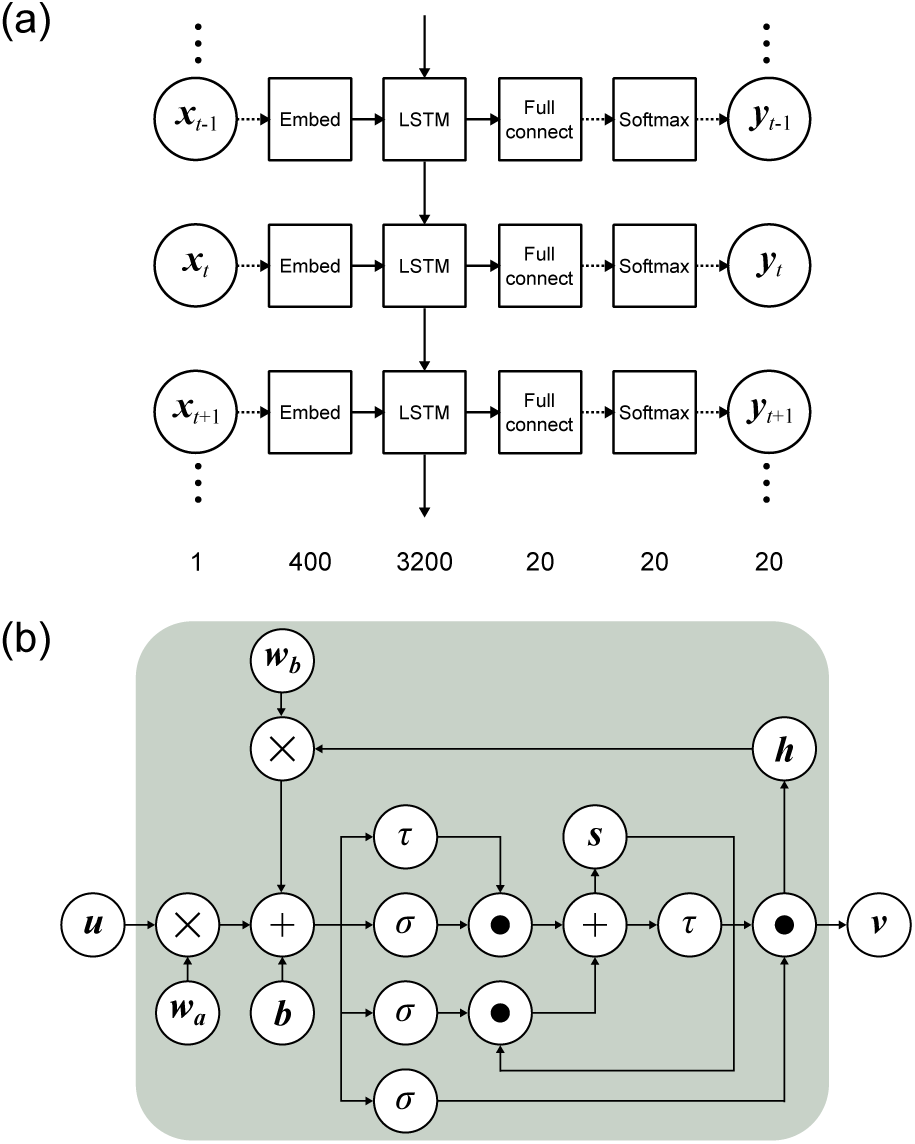
Network of learning. (a) Overview of the designed network in this study. Here, **x**, **y** and *t* represent an input vector, an output vector and a position of an amino acid sequence. In the squares, “Embed”, “Full connect”, “Softmax” stand for a word embedding operation, a fully connected network, and a softmax function layer, respectively. The solid and broken arrows represent a matrix operation and an array operation, respectively. The numbers at the bottom of panel (a) stand for a dimension of vectors of each layer. (b) Description of LSTM layer. Here, **u**, **v**, **h**, **s**, ×, +, dot, *τ*, *σ*, **w_a_**, **w_b_** and **b** stands for an input vector to LSTM unit, an output vector from LSTM unit, a previous input vector, an unit for constant error, a multiplication of matrices, a summation of matrices, a Hadamard product calculation, a hyperbolic tangent, a sigmoid function, a weight matrix to be learned, another weight matrix and a bias vector.

As a framework to implement the learning network, we used Chainer version 1.15.0.1 (Preferred Networks) with CUDA and cuDNN version 6.5 (NVIDIA), and the calculations were performed by a server with Tesla K20m (NVIDIA) at the NIG supercomputer at ROIS National Institute of Genetics in Japan.

### Benchmark of the performance of similarity searches

Performances of profile generators were evaluated based on the results of similarity searches with their generated profiles. As representatives of existing methods of rapid profile generators, we compared our method with CSBuild version 2.2.3 and RPS-BLAST version 2.2.30+. As a test dataset, the SCOP20 test dataset was used, as in the original paper for CSBuild [4], which consists of 5,819 sequences with protein structural information; the maximum percent identity of the sequences in the dataset was less than 20%. In addition to the dataset, we constructed another test dataset as a SCOP20 strict-test dataset. To construct the dataset, we excluded homologous sequences with any sequence in the Pfam40 learning dataset from the SCOP20 test dataset using blastpgp searches with an e-value of less than 10^−5^ as the threshold of homologous hits. As a result, the SCOP20 strict-test dataset contained 1,104 sequences. As a profile library for CSBuild, the data from the discriminative model of CSBuild (K4000.crf) were used. For RPS-BLAST, we excluded all highly similar sequences with any sequence in the SCOP20 test dataset from the conserved domain database for DELTA-BLAST version 3.12 by the same method as that used to make the Pfam40 learning dataset.

To eliminate any biases of alignment algorithms, all profiles in this study were converted to the PSI-BLAST readable format and used as input files in a PSI-BLAST search. As an application of PSI-BLAST, we used blastpgp version 2.2.26 for CSBuild, since CSBuild outputs blastpgp-readable profile files. For the other methods, psiblast version 2.2.30+ was used. There were no significant differences in sensitivity or similarity searchers between these two versions of PSI-BLAST (data not shown). The results of the similarity searches were sorted according to their statistical significance in descending order. Each hit was labeled as a true positive, false positive, or unknown based on the evaluation ruleset for SCOP 1.75 benchmarks [23].. Further, the number of true positives and false positives was normalized by weighting them with the number of members in each SCOP superfamily to negate bias derived from the size of each SCOP superfamily. With this information, we described the receiver operating characteristic (ROC) curves and evaluated the performance [24]. As an evaluation criterion, we used partial area under the ROC curve (pAUC), which is the AUC until one false positive is detected for each query on average. In our case, the pAUC was equivalent to AUC until 1,564 false positives in total were detected, because we weighted detected false positives by the size of each SCOP superfamily, and the number of superfamilies in our test dataset is 1,564.

The profile generation time was benchmarked on an Intel(R) Xeon(R) CPU E5-2680 v2 @2.80 GHz with 64 GB RAM using a single thread.

## RESULTS AND DISCUSSION

### Training a predictor with LSTM

In this study, we assumed profiles generated by HHBlits as ideal profiles and used these as target profiles in training steps. We then attempted to generate profiles as similar to the HHBlits profiles as possible with a predictor using LSTM. The performances of similarity searches with the profiles generated by HHBlits were better than those of the other methods [2].

Initially, we selected amino acid sequences with lengths of 50-1,000 in Pfam40. The sequences did not contain any irregular amino acid characters such as B, Z, J, U, O, or X. We also included 1,329 sequences derived from the SCOP20 learning dataset [4] to the final learning dataset for our reference. As a result, we obtained 1,602,338 sequences and calculated their profiles using HHBlits for each sequence. With this learning dataset, we trained the predictor shown in Figure 1a. For learning, we used 20,000 randomly extracted instances as a validation dataset and checked whether the predictor overfit the training dataset. The number of mini-batches was set to 200, and each amino acid was converted to a 400-dimension float vector by the word embedding method, as described in the methods section. For each sequence, the starting site of learning was not confined to the N-terminal but was selected at random to avoid overfitting of the predictor to the specific site. We observed learning and validation curves to confirm the lack of overfitting and stopped learning at 5,000 epochs (Figure S1). Even using the GPU machine, the completion of our calculations required almost two months.

Using the obtained parameters (weight matrices and bias vectors through the learning), we constructed a novel de novo profile predictor, which we called <u>S</u>ynthetic <u>P</u>rofile <u>Build</u>er (SPBuild). Our profile generator can be downloaded from http://yamada-kd.com/product/spbuild.html.

### Performance comparisons

First, we compared the performance of the similarity searches of the profile generators. The profiles for all sequences in the SCOP20 test dataset were generated by each method, and all-against-all comparisons of the test dataset by PSI-BLAST with the obtained profiles were conducted. As profile generators, we evaluated the de novo profile generators CSBuild and RPS-BLAST, in addition to SPBuild. We also added the performance of PSI-BLAST without iterations (= blastpgp) as a representative sequencesequence-based alignment method for reference. In addition, HHBlits was further compared as another reference, and the results are shown in Figure S2.

As shown in Figure 2a, CSBuild and RPS-BLAST were clearly superior to the sequencesequence-based alignment method, blastpgp. Furthermore, SPBuild showed better performance than those of these methods. When performance was evaluated by the pAUC values (Figures 2b), the values of our method, CSBuild, and RPS-BLAST were 0.217, 0.140, and 0.174, respectively. Notably, the performance of our method (0.217) did not reach that of HHBlits (Figure S2a, pAUC = 0.451), even though we trained our predictor with outputs of HHBlits, indicating that SPBuild was not completely able to mimic the ability of HHBlits. This tendency was also true for another benchmark result, where we evaluated the performance of SPBuild and HHBlits on the SCOP20 learning dataset instead of the test dataset (Figure S2b). Our findings were surprising because the SCOP20 learning dataset was a part of the learning dataset for the construction of the predictor with LSTM, and the performance of our predictor should reach that of HHBlits. One possible reason for the observation is that LSTM may not have worked properly on our learning scheme. To examine this possibility, we performed another learning method to examine the performance of LSTM itself with our learning scheme, where we trained a predictor with only the SCOP20 learning dataset and let the predictor overfit the dataset. As a result, the performance of the predictor was almost the same as that of HHBlits, as expected (Figure S2c). This result indicated that LSTM could precisely learn input sequence properties and output correct PSSMs, but that the performance of the predictor was worse than that of SPBuild with proper learning due to the overfitting of the predictor to the learning dataset (Figure S2d). In short, these results suggested that LSTM worked correctly, and that relationship between performance and overfitting was a simple trade-off. Therefore, we concluded that SPBuild could be trained moderately and pertinently without conflict under our learning dataset and hyperparameters.

**Figure 2.**
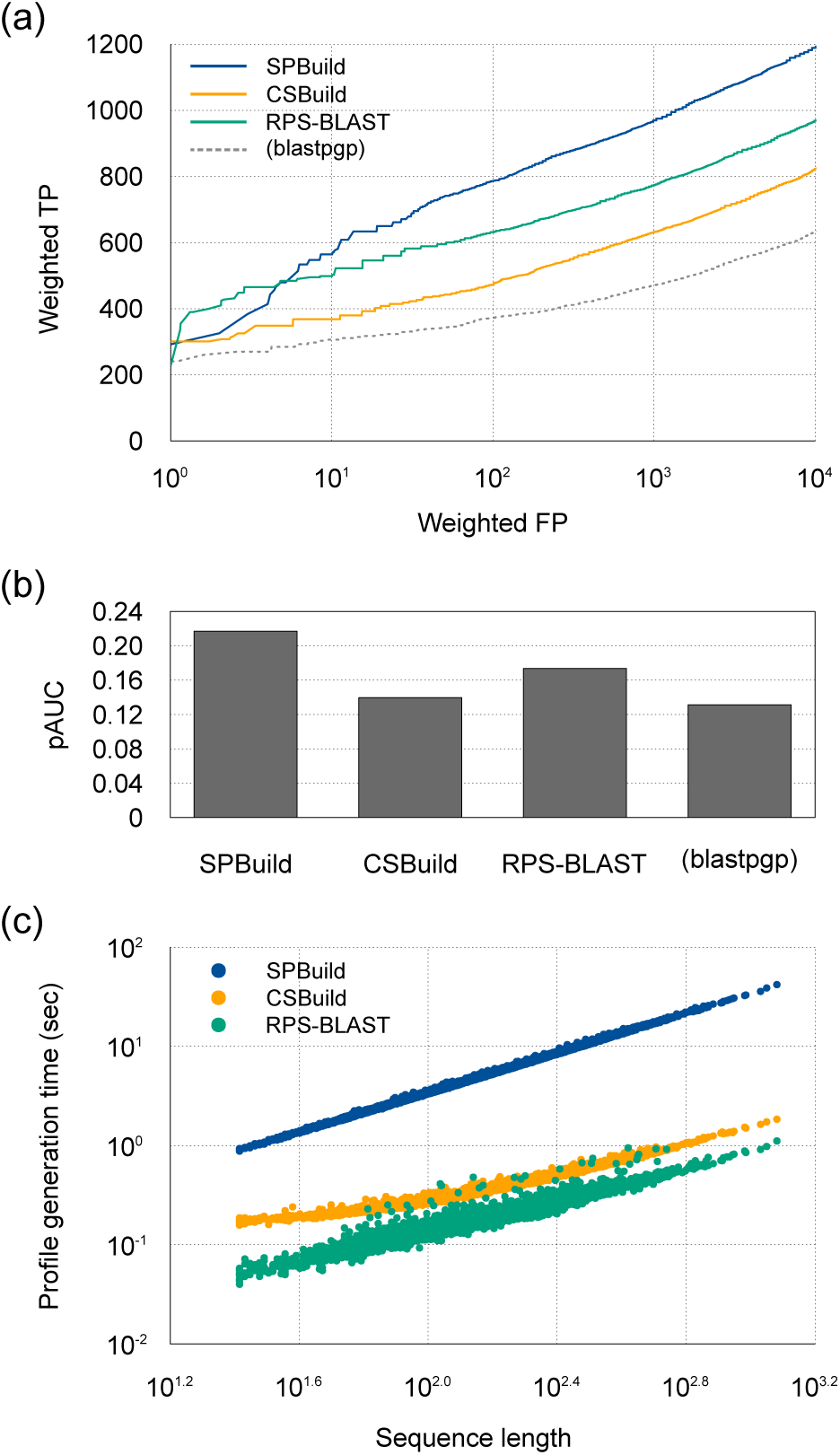
Performance comparisons of (a, b) similarity searches and (c) calculation time. (a)ROC curves of SPBuild and other methods. Here, the performance of blastpgp was added for a reference. (b) AUC 1000 values of SPBuild, CSBuild, RPS-BLAST, and blastpgp. (c) The scatterplot of the profile generation time for each method on the SCOP20 test dataset.

Next, we evaluated the profile generation time of each method. Table1 shows the mean computation time of profile generation using the SCOP20 test dataset. SPBuild was found to be almost 20 times faster than HHBlits, although CSBuild and RPS-BLAST were still faster than SPBuild. However, we think the most important property of a sequence handling method in the big data era is scalability to the data, namely, time complexity of the method against the input sequence length. Theoretically, the time complexity of our method would be linear compared with the input sequence length, similar to CSBuild and RPS-BLAST. To clarify this point, we plotted profile generation times (seconds) versus input sequence lengths (*N*), as shown in Figure 2c. When the instances were fitted to a line, the determination coefficient was 0.998, and the slope of the line was 1.00. This result indicated that the time complexity of our method was *O*(*N*). Notably, the slopes of CSBuild and RPS-BLAST appeared to be less than 1.0 in the figure; however, errors in the experiments or other factors in the implementation of these programs may have caused this because the costs of these calculations must be higher than that of *O*(*N*). Although our method required much time to compute large matrix calculations in the neural network layers and was therefore slower than CSBuild and RPS-BLAST with the currently used sequence database, our method had linear scalability against the number of input sites or sequence length and the number of input sequences.

**Table 1.** Comparison of profile generation times.

|  | Mean | SD |
| --- | --- | --- |
| SPBuild | 5.99 | 3.83 |
| CSBuild | 0.390 | 0.161 |
| RPS-BLAST | 0.208 | 0.102 |
| HHblits | 120 | 105 |

Means and standard deviations (SDs) of profile generation times (s) against 5,819 sequences in the SCOP20 test dataset.

### Memory power of LSTM in our problem

We also examined the memory power of LSTM in our problem in order to determine the feasibility of the LSTM approach for sequence-based predictions. For this purpose, we considered the reset time lengths of memory cells (h in Figure 1b) at sequence lengths of 5, 10, 20, 30, 50, 100, 200, and 300 and for full-length sequences. We then benchmarked the performances of similarity searches with the SCOP20 test dataset. The memory reset time length was directly linked to the memory power of the predictors, and a predictor with a memory reset time length of 5, for example, generated profiles based on information from the previous five sites, including the current site. As a result, the performance of similarity searches clearly changed as the memory power decreased (Figure 3a). We also checked the performance of CSBuild with the same plot (Figure 3a). As described above, CSBuild constructs profiles by merging 13-mer short profiles; thus, we imagined that its performance would be similar to that of the LSTM profile predictors with low memory power. However, we found that the performance of CSBuild was located in the middle between memory powers of 30 and 50 for the LSTM predictors. We are not sure why this happened, but it may be because the sensitivities (corresponding to vertical axis of Figure 3a) of LSTM predictors were worse than expected or because of the excellence of CSBuild implementations.

**Figure 3.**
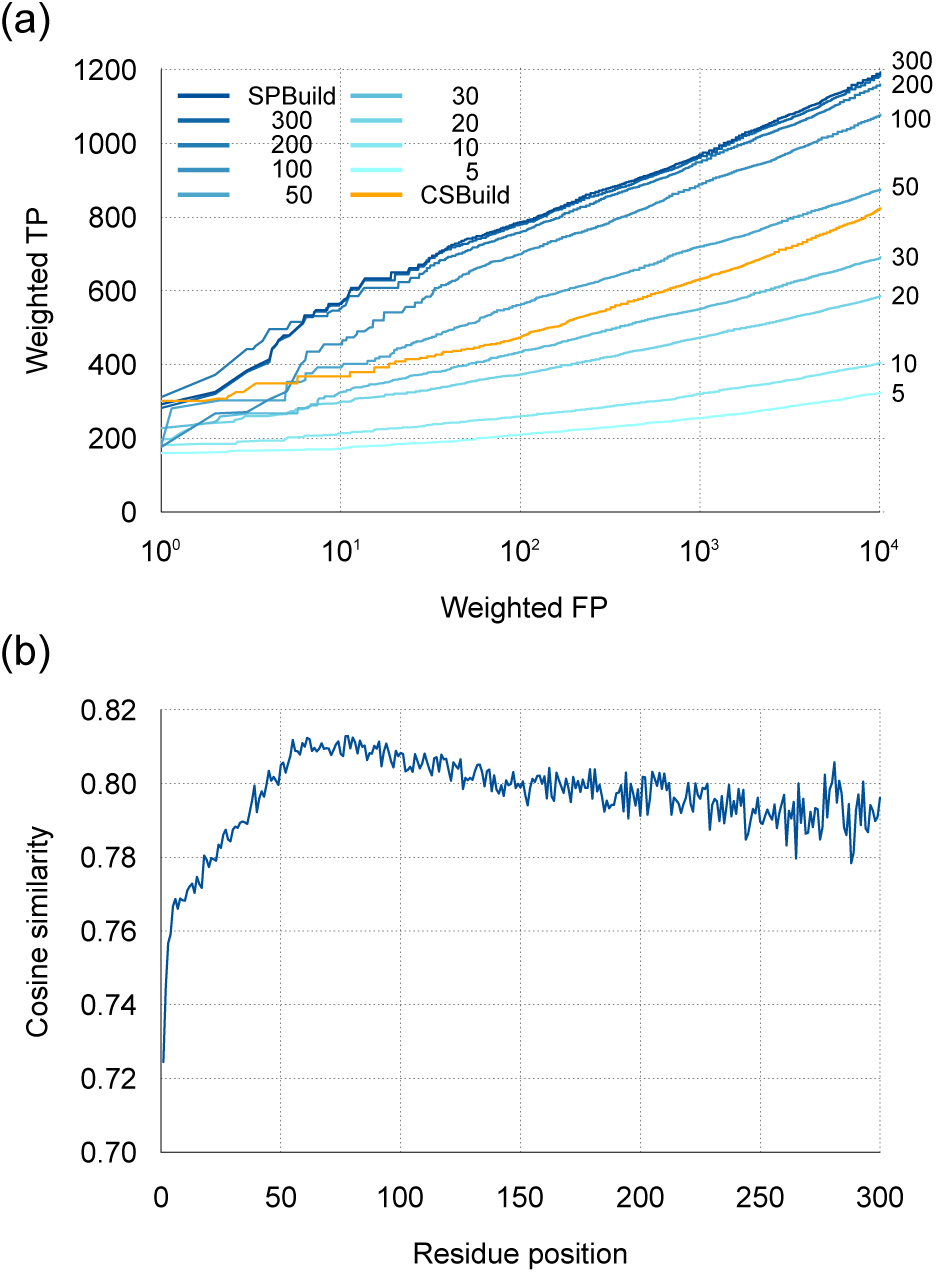
Effects of memory power of LSTM on predictors. (a) Comparison of profile generators with various reset lengths of memory on LSTM. The benchmark dataset was the SCOP20 test dataset. The reset time of SPBuild corresponded to the input sequence length. (b) Mean cosine similarity between output vectors of SPBuild and target vectors as a function of the position of residues in input sequences of the SCOP20 test dataset.

To improve our understanding of the generated profiles by SPBuild, we evaluated the mean prediction accuracy (cosine similarity between output vector, y, and target vector) of SPBuild for each position of a residue on whole input sequences and observed that there was a clear transition in the plot (Figure 3b). The prediction accuracy of the initial portion (~50) was worse than those of the other parts. This lower performance could be caused by the nature of LSTM. LSTM initializes the internal state of memory (h) by a null vector, which does not reflect any features of the learning dataset; thus, the prediction would be not stable until LSTM memorizes and stores a certain level of context information into memory. In our case, the level of context information was 50–60 residues. In addition, the decrease in accuracy in the last part (200) was derived from the nature of our learning dataset; the mean length of SCOP20 was about 154, and SPBuild may be able to be optimized for the average length. This consideration was consistent with the observation that improvement of the performance with memory power of 200 and 300 decreased compared with smaller memory power lengths (Figure 3a). On the basis of the observations that the prediction confidence of the N-terminal region was not good, we think that it might be possible to improve the performance of SPBuild by combining prediction results from both N-terminal and C-terminal directions. Although we did not implement this feature because the learning process took lots of time, this will be a future direction for further improvements.

In conclusion, these results suggested that substantially long length context, ideally speaking, the context of the sequence length of at least more than 50, would be required to predict precise profiles. Protein primary and secondary structures, including solvent accessibility and contact number, must be restricted by protein steric structures, which are formed by complex remote interactions of amino acid residues. Our findings reflect the influence of remote relationships stemming from the steric structure on sequence context. In other words, LSTM will be a powerful predictor for divergent features of proteins, if appropriate memory power length is used. Indeed, other sequence-based predictors using LSTM have achieved successful outcomes and have shown the high feasibility of LSTM [13, 17, 18].

### Long-range interactions and memory lengths

As shown in Figure 4a, we calculated the pAUC values of SPBuild relative to those of CSBuild and RPS-BLAST for each SCOP class. The values were calculated by dividing the pAUC value of SPBuild by that of each method, which indicated how the sensitivity of SPBuild was better than those of the existing methods for each SCOP class. Actual pAUC values are shown in Table S1. Notably, the performance of SPBuild was 2.00- and 1.49-fold higher than those of CSBuild and RPS-BLAST for SCOP class b, respectively. SCOP class b consists of *β* proteins. Generally, *β*-strands are constructed by remote interactions between residues when compared with *α*-helices. Secondary structure predictors with a window-based method developed by machine learning methods tend to show poorer performance in *β*-regions than in α-regions. The main reason for this weakness is related to the long-range interactions in *β* structures, which may not be properly handled by the limited lengths of sequence windows [25, 26]. This tendency may also be observed with the profile predictors. CSBuild constructs final profiles by assembling short window-based profiles, and RPS-BLAST also combines many subjected profiles obtained by local similarity searches against profile libraries. The actual mean length of the profiles evaluated by RPS-BLAST with three iterations (default) on the SCOP20 test dataset was 77, which was relatively longer than that of CSBuild but still shorter than the typical length of a protein. However, our method can theoretically memorize whole-length amino acid sequences and can take the remote relationship into consideration to generate profiles.

**Figure 4.**
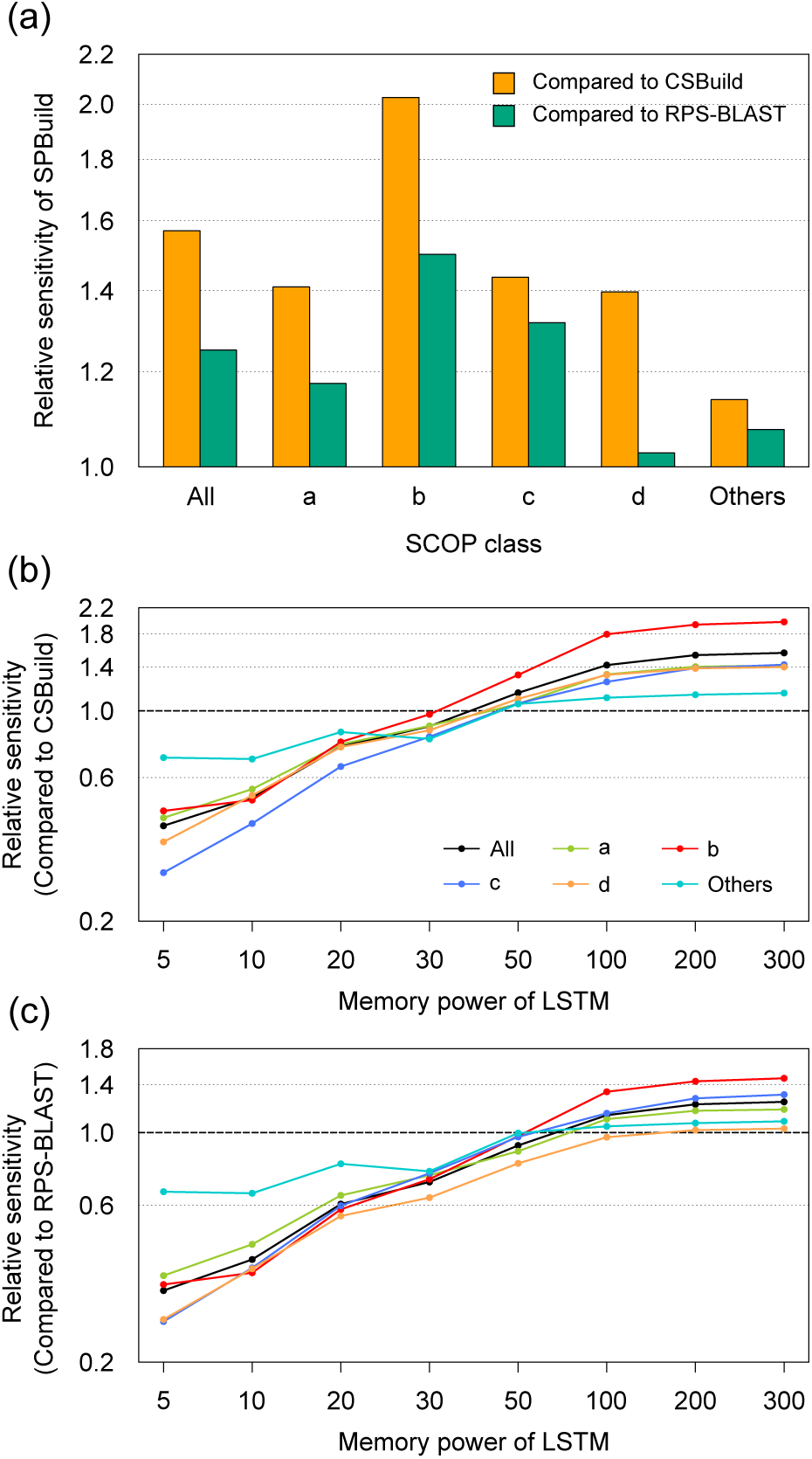
Relative sensitivity of SPBuild against existing methods on the test dataset. (a) The relative sensitivity of SPBuild against existing methods was calculated by dividing the AUC1000 of SPBuild by that of each method. Here, the label ”others” includes SCOP classes e, f, and g. (b) The relative sensitivity of the profile generator with various memory powers of LSTM against CSBuild. (c) The relative sensitivity of the profile generator with various memory powers of LSTM against RPS-BLAST.

To confirm the relationship between memory power length and structural categories, we calculated relative sensitivities for different reset time lengths (Figure 4b and 4c). As a result, the performance improvements in the b category were much better than those of other categories, indicating that memory power was the most important factor for encoding long-range interactions, such as *β* structures.

### Limitation of SPBuild

As described, our method could generate profiles faster than HHBlits and showed higher performance than CSBuild and comparable to or slightly higher performance than RPS-BLAST, particularly for *β* region prediction, possibly due to the memory effects of LSTM. However, there are still some limitations to this method.

One of the limitations of SPBuild is the profile generation time, although the time complexity is linear against input sequence length. SPBuild used huge parameters, particularly for the LSTM layer, to calculate the final profile prediction. Although we set the size of the parameters to the current scale in order to maximize the final performance of SPBuild, we may be able to reduce the size and improve the calculation time if we are able to find more efficient network structures to learn amino acid context. In other words, to resolve the problem, exhaustive optimization of the hyperparameters of LSTM and/or development of novel network structures will be required.

For the construction of the Pfam40 learning dataset, we excluded highly similar sequences with any sequence in the SCOP20 test dataset from the original Pfam40 dataset by blastpgp search having e-value < 10^−10^. It should be noted that the threshold is rather strict to eliminate homologous sequences. In the context of machine learning, the independence of the test and learning dataset is quite important to avoid overtraining, and thus, the same data among the datasets should be eliminated, but similar data are usually retained for better learning. Generally, a test dataset must follow the same probability distribution as that of the learning dataset [27, 28]. In other words, the existence of similar data among a learning and test set is an essential point for supervised learning, and prediction based on supervised learning will fail if no similar data are available among the learning and test dataset. This similar information will be a question of degree, and in our case, better learning would require a homologous relationship in both the learning and test dataset.

Meanwhile, however, in the context of biological sequence analysis, homologous or similar sequences will be conceptual problems. From the viewpoint of machine learning, homologous sequences should not be removed, but conventional approaches of biological sequence analyses usually remove the homologous sequences [29, 30, 31]. For further considerations, we set a moderate e-value threshold of 10-5 aiming to exclude homologous sequences in the Pfam40 learning dataset from the SCOP20 test dataset, and we made another test dataset, a SCOP20 strict-test dataset. According to benchmark results with the dataset (Figure 5), the search sensitivities of de novo profile generators including SPBuild were much lower than that of HHBlits, and our method was worse than blastpgp, which is a sequencesequence-based method. These results will be quite interesting to understand profile generation with machine learning approaches and indicate that machine learning approaches would not be effective at all if homologous sequences are excluded, as conventional sequence analyses methods are doing. In addition, the worse performance of SPBuild might be improved to at least the same level as that of blastpgp by introducing a bailout method, which is a popular approach in machine learning, where profiles are generated from the background frequency of amino acid substitution matrices like BLOSUM [32] or MIQS [33] when the confidences of profile generation are not enough. That kind of bailout is internally implemented by BLAST series, but we did not use it in the current implementations, and thus, it can be a future direction for further improvements.

**Figure 5.**
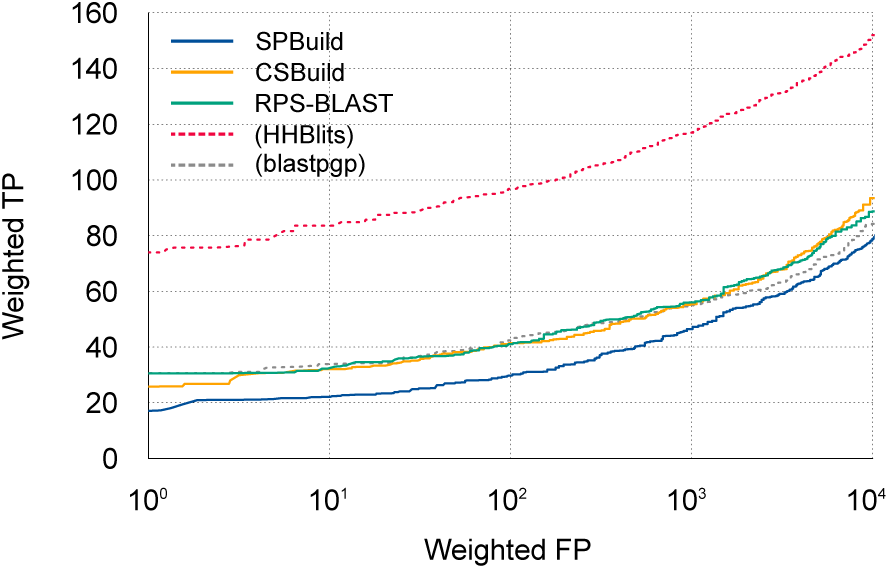
Performance comparisons of similarity searches on SCOP20 strict-test dataset. ROC curves of SPBuild and other methods. The performances of HHBlits (three iterations) and blastpgp were added for a reference.

The performance of iteration search with profiles made by de novo profile generators would be another interesting point for users. To check the performance of iteration searches, we calculated ROC curves for SPBuild, CSBuild, and RPS-BLAST and found that differences in performance became more unclear as the number of iterations increased (Figure S3). The result suggested that the performance of the initial search or qualities of profiles would be of meager importance for the final results in iterative searches if a sufficient number of iterations was used. The reason for this result is unclear; however, we believe that homologous sequences in the sequence space are limited and that almost all homologous sequences can be detected by using modestly good profiles if a large number of iterations are used. Considering the sensitivity of profile sequencebased similarity searches, our method may not be too attractive; however, there are many other uses for profiles. For example, profileprofile similarity searches, where profiles are generated by iterative searches of whole datasets, will be candidates for the application of our approach. The bottleneck of profileprofile searches may be easily resolved with the rapid profile generator. In addition, profiles are often used to encode amino acids into input vectors in other machine learning methods. Machine learning methods generally require large learning data, and currently, long-time iterative searches should be avoided because the calculation time increases depending on the learning data size. In such cases, higher speeds and accurate profile generators will be quite useful.

## CONCLUSION

In this study, we developed a novel de novo generator of PSSMs using a deep learning algorithm, the LSTM network. Our method, SPBuild, improved the performance of homology detection with a more rapid computation time than that of existing de novo generators. However, our goal was not to just provide an alternative method for profile generators but also to elucidate the importance of sequence context and the feasibility of LSTM for overcoming the sequence-specific problem. Our analyses demonstrated the effectiveness of memories in LSTM and showed that SPBuild achieved higher performance, particularly for *β*-region profile generation, which was difficult to predict by window-based prediction methods. This performance could be explained by the fact that our method utilized the LSTM network, which could capture remote relationships in sequences. Moreover, further analyses suggested that substantially long context was required for correct profile generation. We also reconfirmed several limitations of deep learning on our problems. For example, the deep architecture to realize higher performance required considerable computation time, and the intensive elimination of homologous information between the learning and test dataset might make the inference by deep learning impossible. These findings may be useful for the development of other prediction methods.

Profiles are the most fundamental data structures and are used for various sequence analyses in bioinformatics studies. Using SPBuild, the performance of sophisticated comparison algorithms, such as profileprofile comparison methods and multiple sequence alignment, can be further improved. In addition, profiles generated by SPBuild can be useful as input vectors for other machine-based meta-predictors of protein properties.

## ADDITIONAL INFORMATION

### Acknowledgements

We are grateful to Kentaro Tomii and Toshiyuki Oda for constructive discussion. Computations were partially performed on the NIG supercomputer at ROIS National Institute of Genetics and the supercomputer system Shirokane at Human Genome Center, Institute of Medical Science, University of Tokyo.

### Availability of data and material

The source code of SPBuild are available at http://yamada-kd.com/product/spbuild.html.

### Abbreviations

HMM: hidden Markov model; LSTM: long short-term memory; pAUC: partial area under the ROC curve; PSSM: position-specific scoring matrix; ROC: receiver operating characteristic; RNN: recurrent neural network Competing interests The authors declare that they have no competing interests.

